# Evolution of chalcone isomerase from a non-catalytic ancestor

**DOI:** 10.1101/174128

**Authors:** Miriam Kaltenbach, Jason R. Burke, Mirco Dindo, Anna Pabis, Fabian S. Munsberg, Avigayel Rabin, Shina C. L. Kamerlin, Joseph P. Noel, Dan S. Tawfik

## Abstract

The emergence of catalysis in a non-catalytic protein scaffold is a rare, unexplored event. Chalcone isomerase (CHI), a key enzyme in plant flavonoid biosynthesis, is presumed to have evolved from a non-enzymatic ancestor related to the widely-distributed fatty-acid binding proteins (FAPs) and a plant protein family with no isomerase activity (CHILs for “CHI-like”). Ancestral inference confirmed that CHI evolved from a protein lacking isomerase activity. We also identified four alternative founder mutations, *i.e.* mutations that individually instated activity, including a mutation that is not phylogenetically traceable. Despite strong epistasis in other cases of protein evolution, CHI’s laboratory reconstructed mutational trajectory shows weak epistasis. Thus, enantioselective CHI activity can readily emerge despite a catalytically inactive starting point. X-ray crystallography, NMR, and MD simulations reveal reshaping of the active site toward a productive substrate-binding mode and repositioning of the catalytic arginine that was inherited from the ancestral fatty-acid binding proteins.

## Introduction

Enzymes generally diverge from other enzymes by exploiting preexisting promiscuous activities while maintaining key catalytic residues. Moreover, enzymes often evolve into regulatory or scaffolding proteins through loss of catalytic residues^1^. However, the opposite evolutionary trend seems exceedingly rare - emergence of enzyme catalysis starting from a non-catalytic, ancestral protein^2–5^.

Chalcone isomerase (CHI) is enigmatic in this regard. CHI is a key enzyme in the biosynthesis of plant flavonoids – specialized metabolites involved in diverse biotic and abiotic functions including UV protection, flower color, pollen development, root nodulation, plant architecture and chemical defenses^6^. CHI catalyzes the enantioselective formation of the tricyclic flavanone (*S*)-naringenin from its bicyclic precursor chalconaringenin (**Fig. 1a**). However, the only two homologous protein families identified to date are non-enzymatic^7^: the phylogenetically dispersed and presumably more ancient, fatty-acid binding proteins (FAP), and a family of closely related plant proteins dubbed CHI-like (CHIL). FAPs play a role in fatty-acid biosynthesis ^7^ and are found in plants as well as in some algae, protists and bacteria. In contrast, CHI and CHIL have been identified only in plants. CHILs exhibit no CHI activity. Their expression correlates with flavonoid biosynthesis^8,9^, but their biochemical activity remains unknown. Given that neither FAPs nor CHILs show CHI activity, and that FAPs are likely the oldest of the three families, the parsimonious hypothesis is that CHI evolved from a non-enzymatic protein^7^. However, given the evolutionary distance between FAPs and CHIs, and the rarity of cases where enzymes evolved from non-catalytic proteins, CHI’s ancestor may have been an isomerase, and activity was subsequently lost in CHILs. Phylogeny alone cannot reveal when CHI catalysis emerged, nor the trajectory that led from a non-enzymatic ancestor to a bona fide CHI. Therefore, we set out to address these questions using a combination of ancestral sequence inference, directed evolution, x-ray crystallography, NMR, and MD simulations.

**Figure 1.**
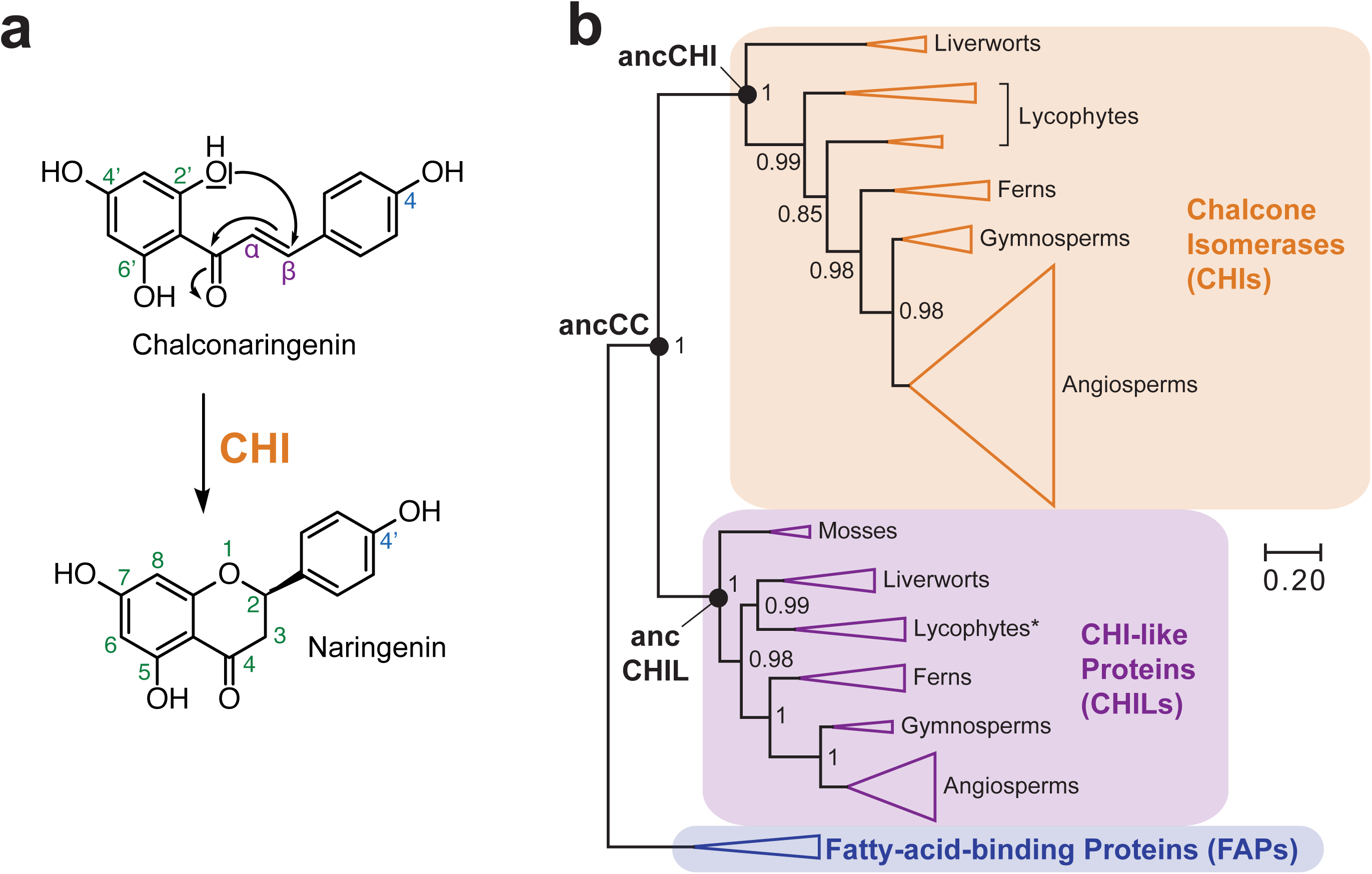
The CHI protein family. (**a**) CHI catalyzes the intramolecular Michael addition of chalconaringenin (2’,4,4’,6’-tetrahydroxychalcone or (*2E*)-3-(4-hydroxyphenyl)-1-(2,4,6-trihydroxyphenyl)prop-2-en-1-one) to yield the flavonoid naringenin (5,7-Dihydroxy-2-(4-hydroxyphenyl)chroman-4-one). (**b**) Cartoon representation of CHI’s phylogenetic tree. Posterior probabilities and the inferred ancestral nodes are annotated. The complete phylogenetic tree is shown in **Supplementary Fig. 2** and the alignment used to generate it is given in **Supplementary Fig. 1** and **Supplementary Data Set 1**.

## Results

### Ancestral sequence inference

We inferred three ancestral nodes by maximum-likelihood: The most probable ancestor of all chalcone isomerases (ancCHI), of all CHI-like proteins (ancCHIL), and the CHI/CHIL common ancestor (ancCC). Given the wide divergence between CHIs and FAPs, a very earlier ancestor was not inferred. Details of the procedure and prediction statistics are provided in **Supplementary Results** and **Online Method**s. Briefly, because protein sequence divergence between CHI, CHIL, and FAPs is high, and includes insertions and deletions (InDels), we generated a structure-based alignment (**Supplementary Table 1, Supplementary Fig. 1**, **Supplementary Data Set 1**). No systematic InDels were found between the CHI and CHIL lineages. Hence, the structural alignment was trimmed in loop regions and at the N- and C-termini, and a phylogenetic tree was generated (**Fig. 1b**; see **Supplementary Fig. 2** for the complete tree). Remaining gaps and ambiguously aligned positions were handled manually in the reconstructed ancestors (**Supplementary Table 2**, **Supplementary Fig. 3**).

Five residues were previously identified as important for catalysis in extant CHIs^7,10^ and, as expected, these were all present in the most probable ancCHI sequence (posterior probabilities p = 0.84 – 0.98). Of these, R34, T46, Y104 (ancCC sequence numbering is used throughout), which are highly conserved in extant CHIs (80-100%, **Supplementary Fig. 1**, **Supplementary Data Set 1**), are present in ancCC (p = 0.97-0.98). For the remaining two positions, different amino acids were predicted in ancCC vs. ancCHI (S111 vs. N and V188 vs. T). These residues were shown to be functionally less relevant^10^ and accordingly more variable (67-68%) in extant CHIs, as reflected by lower posterior probabilities in ancCHI (0.84 and 0.91).

### The inferred CHI/CHIL ancestor exhibits no CHI activity

The three inferred ancestors expressed in high yield as soluble proteins in *E. coli* and displayed high thermal stability (apparent midpoint melting temperatures (T_m_) > 75 °C; **Supplementary Table 3**, **Supplementary Fig. 4**). AncCHI was an active CHI with a *k*_*cat*_/*K*_*M*_ of 2.1 × 10^5^ M^-1^s^-1^ for (*S*)-naringenin formation, ∼40-fold lower than that of *Arabidopsis thaliana* CHI (*At*CHI, 7.7 × 10^6^ M^-1^s^-1^, **Fig. 2a**; **Supplementary Table 4**). Like all characterized extant CHIs, ancCHI enantioselectively produces only (*S*)-naringenin (**Fig. 2b**; **Supplementary Table 4**). Isomerase activity above background was not detected either in ancCHIL (as in extant CHILs) or in ancCC (**Fig. 2a**, **Supplementary Fig. 5**). Nonetheless, as detailed below, the crystal structure of ancCC showed the characteristic CHI-CHIL fold. Therefore, ancCC appears to be correctly folded and stable, suggesting that the lack of enzymatic activity is not due to inference errors. However, we only reconstructed the statistically most probable sequence while in effect, ancestral inference yields a “cloud” of sequences that relate to the historical ancestor^11–13^. Results described below indicate that this “cloud” includes sequences that possess basal CHI activity, but the probability that ancCC’s isomerase activity exceeded low, promiscuous levels is negligible. Therefore, our findings confirm the hypothesis that CHI evolved from a non-enzymatic protein.^7^

**Figure 2.**
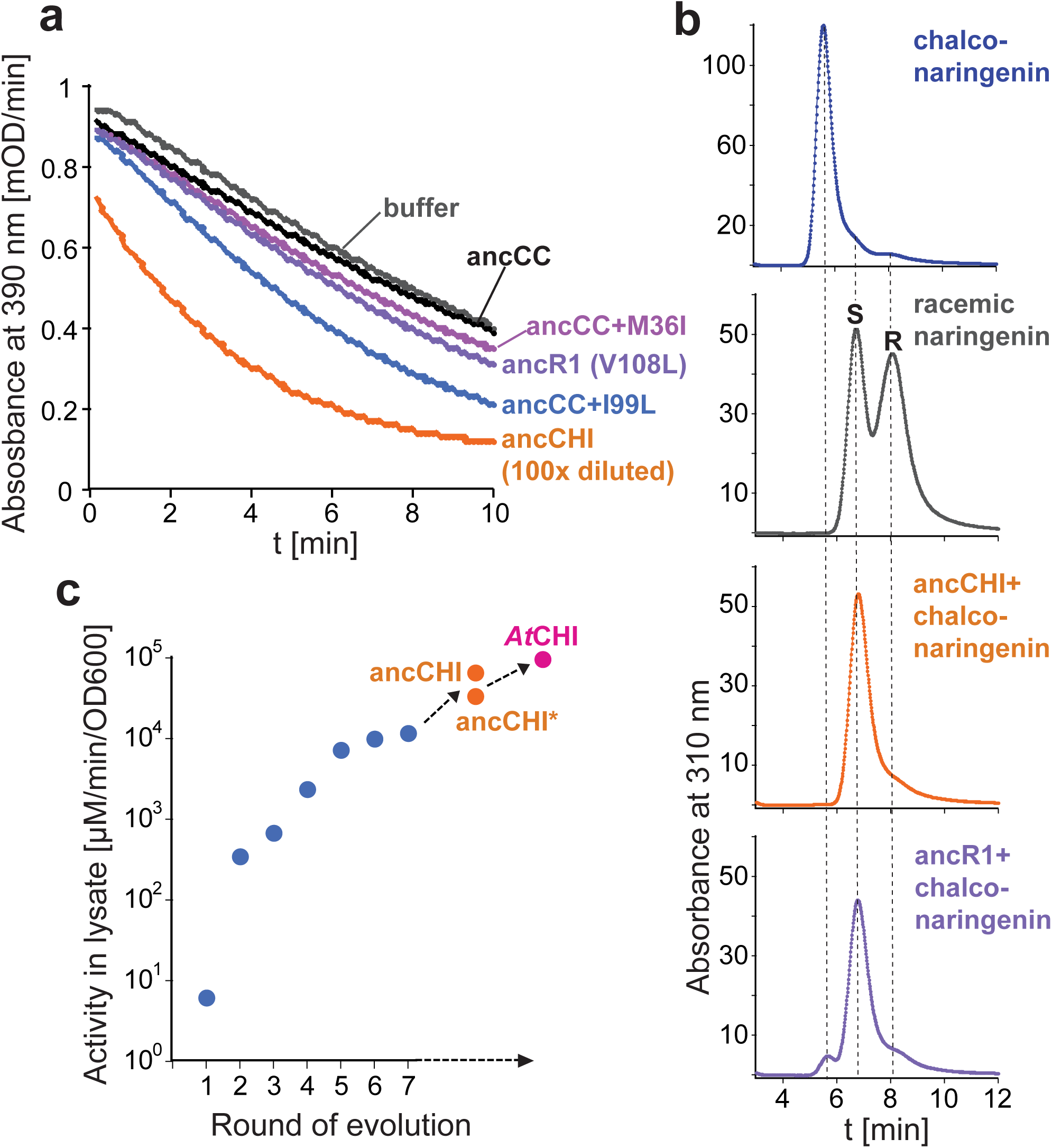
Isomerase activity of the inferred ancestors and evolutionary intermediates. (**a**) Kinetics of naringenin formation indicated by the decrease in 390 nm absorbance. While ancCHI is a highly active isomerase, no turnover of naringenin chalcone above background (buffer) could be detected for ancCC at ≥100-fold higher protein concentrations. The three founder mutations induce a low yet reproducible rate enhancement. (**b**) Steroselectivity of CHI variants. Like all characterized extant enzymes, all evolutionary intermediates produce (S)-naringenin. (**c**) The evolutionary trajectory leading from ancCC to ancCHI*. The underlying mutatons are shown in **Supplementary Table 8**. The Y-axis denotes isomerase activity under the screening conditions (*i.e.*, in clarified lysates of the E. coli cells in which these variants were expressed). AncCHI and AtCHI are shown for comparison. Specific activities measured with purified proteins and kinetic parameters are provided in **Supplementary Tables 4** and **9**.

### An adaptive trajectory from an inactive ancestor to active CHI

Can a putative, historically relevant mutational trajectory be reconstructed that connects ancCC and ancCHI? We addressed this question using a phylogenetic library approach, *i.e.* by examining various combinations of the amino acid exchanges that separate the inferred ancCC and ancCHI sequences. While these two sequences differ by 65 amino acids (out of 220), we posit that most of these non-conserved exchanges have no significant contribution to the acquisition of enzymatic activity. Therefore, to limit library size, we constructed a variant of ancCHI, dubbed ancCHI*, where exchanges in positions that are non-conserved or/and distant from the active site were excluded. AncCHI* differed from ancCC at 39 positions, possessed an identical structure (elaborated below), and exhibited a *k*_*cat*_/*K*_*M*_ of 1.0 × 10^5^ M^-1^s^-1^ (∼2-fold lower than ancCHI; **Supplementary Fig. 6**; **Supplementary Table 4**). The first library was generated by randomly incorporating 39 oligonucleotides into the ancCC gene, each encoding one amino acid exchange, at an average rate of 2 exchanges per gene (**Supplementary Tables 5-6**). Expression in *E. coli* and activity-based screening of >700 clones in clarified cell lysates identified several variants with isomerization rates above background (**Fig. 2a**). Several mutations were enriched, in particular L108V (15 out of 23 clones, **Supplementary Table 7a**). A variant containing only L108V was also identified (∼2.5-fold increased isomerization rate compared to the uncatalyzed background reaction). The L108V variant, hereafter called ancR1, was purified and its catalytic activity verified (**Fig. 2b**; **Supplementary Tables 4** and **8**). The activity of this and other early mutants was too low to accurately determine *k*_*cat*_ and *K*_*M*_ values, but assays with purified proteins consistently indicated rate enhancements above background in agreement with the lysate measurements (**Supplementary Tables 4** and **9**).

Six additional rounds of library construction (alternating between spiking of ancestral mutations and shuffling of improved variants) and screening were performed, yielding variant ancR7 that exhibited only 3-fold lower catalytic efficiency compared to ancCHI* (*k*_*cat*_/*K*_*M*_ = 7.6 × 10^4^ M^-1^s^-1^; **Supplementary Tables 4, 6-8**). All variants showed the same enantioselectivity as extant CHIs (**Fig. 2b**; **Supplementary Table 4**), suggesting that CHI’s enantioselectivity was embedded in the inactive ancCC scaffold. The emerging evolutionary trajectory represents a viable adaptive pathway with gradual increases in activity and diminishing returns in later rounds (**Fig. 2c**)^14^. Interestingly, ancR7 contained only 4 of the 5 previously identified catalytic amino acids (S111N fixated, but not V188T) indicating that the V188T exchange plays a relatively minor role in CHI’s catalytic emergence.

### CHI activity can arise independently via different non-epistatic founder mutations

Our initial screen identified L108V as the likeliest founder mutation, *i.e.* a single mutation that can initiate CHI activity on its own. To identify alternative founder mutations, we screened a second R1 library with a low mutation rate (0.2 amino acid exchanges/gene) such that all possible single mutants were likely represented (>900 clones, ∼5× oversampling, **Supplementary Table 6**). Two additional founder mutations were identified (**Fig. 1a**): M36I, resulting in a 1.3-fold higher catalytic rate compared to the background reaction, and I99L with a 2.0-fold increase. To explore possible second mutations, we screened three low-mutation rate R2 libraries, each based on one of the three founder mutations (**Supplementary Table 6**). The improved variants each contained the remaining founder mutations (**Supplementary Tables 10a-d**). Indeed, the effect of these 3 founder mutations is largely additive since all possible combinations of these mutations gave a smooth, ‘uphill’ trajectory (**Fig. 3a**). The absence of strong epistasis is further evidenced by the fact that reversion of any of the three founder mutations in ancCHI* resulted in only minor reductions in *k*_*cat*_/*K*_*M*_ (4-6-fold, **Supplementary Table 9**).

**Figure 3.**
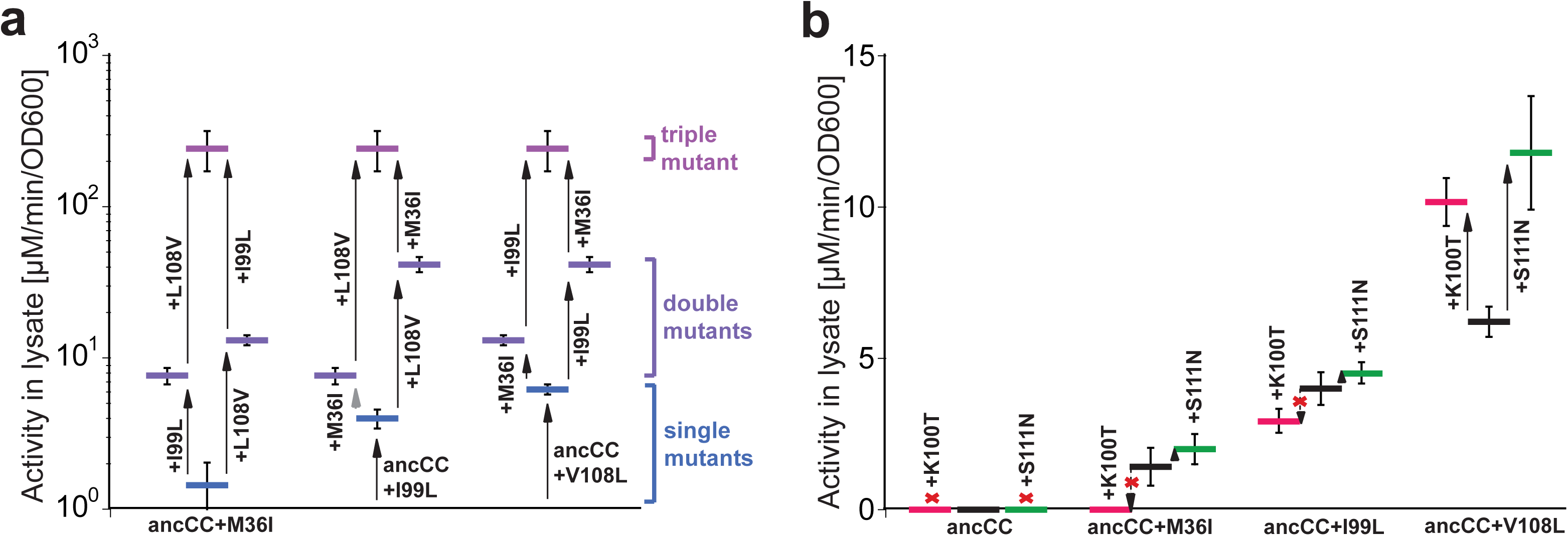
Additivity versus epistasis along CHI’s evolution. (**a**) The three founder mutations (L108V, I99L and M36I) can be combined in any order in an uphill trajectory. (**b**) Two additional mutations that appeared later in the trajectory, K100T and S111N, only develop their effect if L108V is present. Initial rates were determined in clarified cell lysates. Specific activities obtained with purified protein are provided in **Supplementary Tables 4** and **9**.

In addition to various combinations of founder mutations, the L108V library contained improved variants with either K100T or S111N as the second mutation, the latter being one of the two remaining active-site exchanges discussed above. A combinatorial mutational analysis confirmed that neither of these second mutations can serve as a founder mutation, as on their own they do not elicit activity in ancCC (**Fig. 3b**). This analysis also explained why neither K100T nor S111N were found in any selected variant: in the background of either M36I or I99L, K100T was nearly neutral, while S111N reduced activity. That the effects of K100T and S111N develop only in the presence of L108V reinforces the key role of this mutation in initiating CHI’s catalytic emergence.

By the fourth round, all three founder mutations, M36I, I99L, and L108V, and both second-stage mutations, K100T and S111N, were fixed, and together, accounted for most of the activity gains. The quintuple mutant is only 4-fold less active in lysate and 5-fold in terms of *k*_*cat*_/*K*_*M*_ compared to ancR4, which contains six additional mutations.

### An alternative pathway not seen in CHI’s phylogenetic history

The above experiment sampled mutations that are phylogenetically traceable, *i.e.*, sequence exchanges between the inferred ancestral nodes ancCC and ancCHI. However, alternative routes may exist that were either not explored by natural evolution or were historically explored, but are no longer visible in the phylogenetic record. We thus subjected ancCC to random mutagenesis. We performed four rounds of random mutagenesis by alternating between error-prone PCR (1.5 amino acid substitutions/gene on average) and DNA shuffling, and screening for CHI activity (**Fig. 4a**; **Supplementary Tables 6, 11**). Approximately 1100 clones were sampled in round 1. Strikingly, none of the above-mentioned phylogenetic founder mutations was observed. Most of the active variants identified were mutated at position F133 – predominantly to Leu (7/19 sequenced clones) and also Ile or Ser (2 clones each). Thus, F133L seems like an alternative to the phylogenetic trajectory, as far as this trajectory can be inferred from the extant sequences. While there is a reasonable probability that Leu was present in ancCC (p(L)=0.12 compared to p(F)=0.84), it is negligibly predicted in ancCHI (p(F)=0.90, p(L)<0.01, **Supplementary Table 12**). Accordingly, position 133 is predominantly Phe in extant CHIs (85%) while Leu is rare (3.4%) and Ile and Ser are not found (**Supplementary Table 12**). By round ep4, higher activity evolved and four additional mutations fixated (**Fig. 4a**, **Supplementary Table 13**), three of which resemble F133L in not being widely represented in extant CHIs and not exhibiting significant prediction probabilities in ancCHI (**Supplementary Table 12**). Nonetheless, as observed in the ancestral trajectory, all variants exclusively produced (*S*)-naringenin (**Supplementary Table 4**), reinforcing our hypothesis that enantioselectivity was embedded in ancCC’s pre-catalytic site.

**Figure 4.**
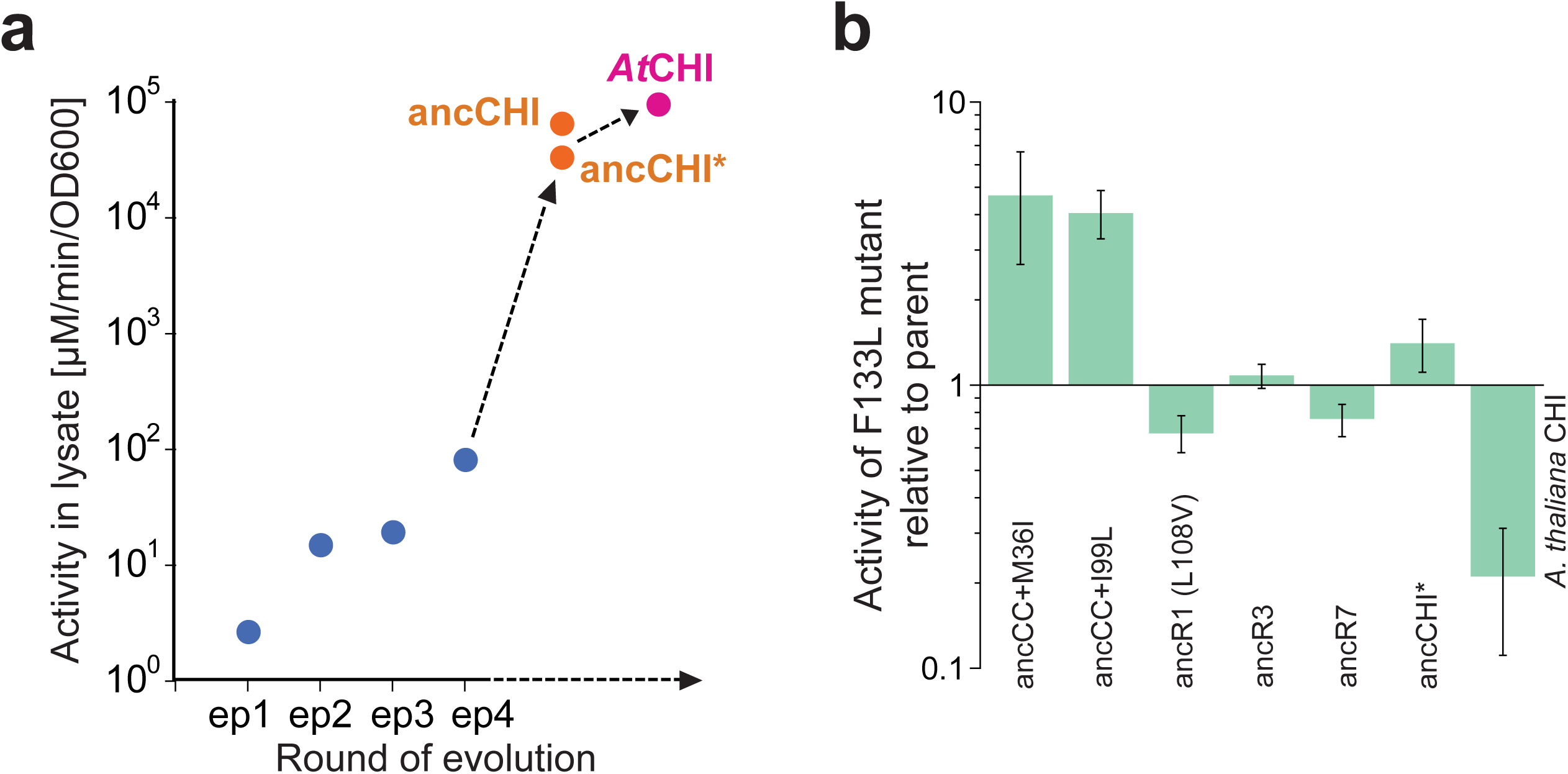
The alternative random mutagenesis trajectory. (**a**) Development of isomerase activity along the trajectory (as in **Fig. 2c**). The underlying mutations are shown in **Supplementary Table 13**. (**b**) The effect of F133L, the founder mutation of the alternative trajectory, changes depending on the genetic background. Shown is the ratio of activities of different variants with and without the F133L mutation. Specific activities and kinetic parameters measured with purified protein are provided in **Supplementary Tables 4** and **9**. F133 corresponds to F146 in *At*CHI.

At first glance, it may seem like the error-prone trajectory, which reached a *k*_*cat*_/*K*_*M*_ of only 1.6 × 10^3^ M^-1^s^-1^ (**Fig. 4a**, **Supplementary Table 4**), lags behind the ancestral trajectory (**Fig. 2c**). However, the two trajectories are not directly comparable. Firstly, the ancestral libraries primarily contained beneficial or neutral mutations, while the vast majority of random mutations are deleterious^15^. Secondly, ancR4 contains 11 mutations compared to only five in epR4. However, the key question is whether the two trajectories are bridgeable, or incompatible as commonly observed in evolutionary trajectories (*i.e.*, combining their mutations leads to loss of activity)^16–18^. To this end, we introduced the random founder mutation F133L into various ancestral intermediates (**Fig. 4b**, **Supplementary Table 9**). F133L is slightly deleterious in the background of the most catalytically active ancestral founder mutation (L108V) yet beneficial in the two others (M36I, I99L). When added to the more advanced intermediates (R3, R7 and even ancCHI*), F133L exerts no significant effect, although it significantly reduces the activity of *At*CHI. Thus, an alternative trajectory based on F133L is in fact compatible with the phylogenetic trajectory. That F133L is deleterious in an extant CHI indicates that incompatibility is the outcome of the later stages of evolution, and also explains why Leu at position 133 is inferred with negligible probability in CHI’s ancestor.

Overall, the above results reinforce the conclusion of facile emergence of CHI despite its origin from a catalytically inactive ancestor - multiple founder mutations and subsequent trajectories are available with unexpectedly weak functional epistasis. In other words, the evolutionary landscape underlying CHI’s emergence is smooth rather than rugged.

### The structural and mechanistic basis of CHI’s emergence

CHI’s catalytic activity has been attributed to five residues^7,10,19^. R34 aligns the substrate and stabilizes the transition-state by interacting with the attacking 2’-hydroxyl (**Fig. 1a**, **Fig. 5a**)^20^. The other four residues form hydrogen bonds with the carbonyl oxygen (T46, Y104) and 6’-OH (N111, T188; ancCC numbering) thus directing substrate binding and favoring ring closure to give the *(S)*-enantiomer (**Fig. 5a**). R34, T46, and Y104 are most critical because their mutagenesis leads to dramatic losses in activity (100-1000 fold reduction in *k*_*cat*_ ^10,20^) and these three residues are already present in ancCC (**Fig. 5b**). The catalytic arginine dates back to FAPs, where it is absolutely conserved and coordinates the negatively charged carboxylate of the bound fatty acids^7^. Why then is the inferred ancestor catalytically inactive, and how do the identified mutations enable the emergence of catalysis and enantioselectivity?

**Figure 5.**
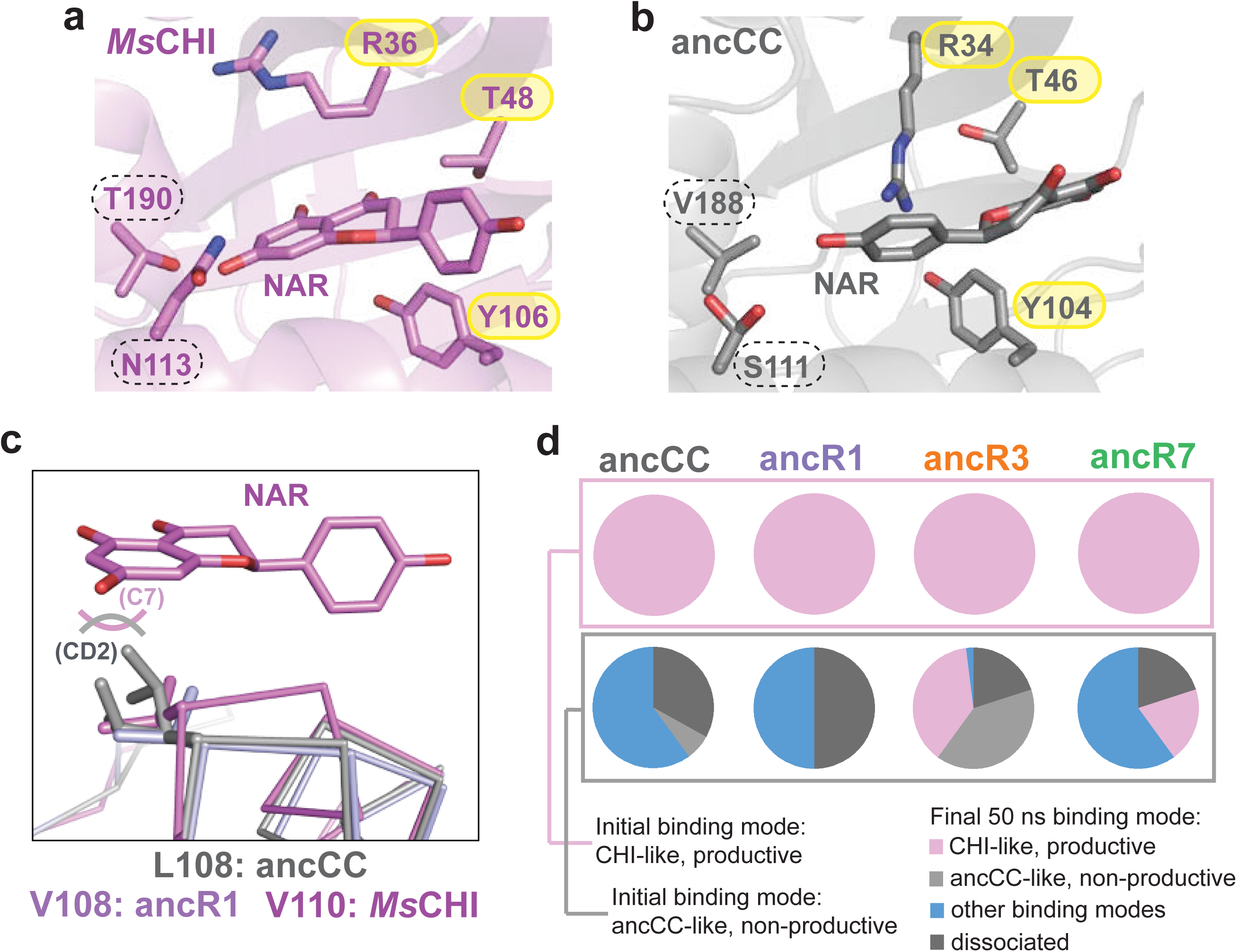
Structural changes over the evolution. (**a**) Crystal structure of *Ms*CHI [19] with (*S*)-naringenin (NAR) bound in the CHI-like mode. Selected active-site residues are labelled. (**b**) Crystal structure of ancCC with naringenin bound in the ancCC-like mode. The three key residues are already present (yellow circles). (**c**) The structural alignment of ancCC and ancR1 with *Ms*CHI with (*S*)-naringenin bound (Ca alignment with PyMOL, RMSD = 1.14 Å) reveals a likely steric clash between (*S*)-naringenin and the side chain of L108 in ancCC (distance CD2 to C7 ∼2 Å), suggesting that the CHI-like binding mode is incompatible with ancCC. AncR1 harbors an activating substitution, L108V, which alleviates the steric clash by reducing the length of the side chain by a methylene moiety, thus increasing active site volume. (**d**) Percentage of simulation time in which the substrate assumes a particular binding mode in ancCC, ancR1, ancR3, and ancR7 after initially being placed in either the productive (top panel) or non-productive mode (bottom panel). Each variant was simulated in five replicas (100 ns each). The last 50 ns of each simulation were combined and RMSD-based clustering using the average linkage algorithm was applied to calculate the fraction of time in which the substrate resides in the productive, non-productive, or other binding modes (illustrated in **Supplementary Fig. 11**) or dissociates from the active site. Note that productive and non-productive modes include structures in which the substrate is not ideally aligned with either of the two modes, with some RMSD fluctuations still observed, but retaining the proper orientation and being fully bound in the active site. Additional simulations performed with ancCHI and *At*CHI are shown in **Supplementary Fig.s 9-10**.

We solved the x-ray crystal structures of various evolutionary intermediates (**Supplementary Table 14**, **Supplementary Fig. 7**). The observed changes are subtle and relate to the repositioning of specific active-site side chains rather than to global rearrangements (**Supplementary Table 15**). Along the trajectory, mutations occurred from the inside out, *i.e.* starting with first-shell mutations and progressively outward from the active site to 2^nd^ and 3^rd^ shells (**Supplementary Fig. 8**). This trend suggests that the late mutations fine-tune the structural effects of the early mutations^14^.

Most noticeably, substitution of L108 by the shorter Val side chain in ancR1 enlarges the active-site cavity thus favoring productive substrate binding modes (**Fig. 5c**). When crystals of ancCC (1.5 Å resolution) and ancR1 (1.4 Å) were soaked with racemic (*R/S*)-naringenin (**Fig. 5b**, **Supplementary Fig. 7**), we observed a ligand orientation that is the opposite of the one observed in *Medicago sativa* CHI (*Ms*CHI, PDB ID 1EYQ, **Fig. 5a**)^19^ and likely is non-productive. Because complexes could not be obtained for the other variants, we performed MD simulations to further explore the two contrasting binding modes (dubbed “ancCC-like” and “CHI-like”). Specifically, we computationally placed chalconaringenin in either the productive or non-productive orientation into the crystal structures of various evolutionary intermediates, and followed their positional changes over time (**Fig. 5d**, **Supplementary Figs. 9-10**). When the substrate was placed in the CHI-like mode, it remained in this orientation in all structures, supporting that this mode is indeed the productive one (**Fig. 5d**). However, when the substrate was initially placed in the non-productive ancCC-like mode, a complex trend was observed. In both ancCC and ancR1, the substrate tends to either dissociate or remain in the active site while adopting a variety of alternative non-productive orientations (**Fig. 5d**, **Supplementary Fig. 11**). In ancR3, a more advanced evolutionary intermediate, productive binding was first observed and in ancR7, non-productive ancCC-like binding was eliminated (**Fig. 5d**). Control simulations of the ancCC and ancR1 crystal structures in complex with (*S*)-naringenin showed that the product complexes were stable over the course of the simulation, indicating that the observed instabilities of the ancCC-like substrate binding mode are not a simulation artifact (**Supplementary Fig. 12**). Thus, one important factor in the emergence of CHI activity is a change in substrate positioning.

Concomitantly, changes in the conformational ensemble of the catalytic arginine occurred. R34 assumes various rotamers in the structures of different evolutionary intermediates, and also in extant CHIs (overall, the position of the guanidinium carbon varies by up to 7.7 Å; **Fig. 6a**). Nonetheless, it appears that a consistent change in the most populated rotamers accompanied the emergence of isomerase activity, as indicated by both NMR and MD simulations. The NMR chemical shift of the guanidinium N^ε^ is well resolved. Three other arginines that reside on the protein surface were mutated to lysines (R113K, R115K, R134K) in key evolutionary intermediates (ancCC-, ancR1-, ancR3- and ancR7-ΔR; see Online Methods). This resulted in an unambiguous resonance signature for the catalytic arginine side chain in the ^15^N-^1^H HSQC spectra of the studied variants (**Supplementary Fig. 13**). Aggregation of ancR3-ΔR excluded it from our NMR analyses, but changes in the steady-state heteronuclear NOE effect were readily measured in the other variants, providing dynamics of pico- to nanosecond thermal fluctuations of ^15^N-^1^H bond vectors^21^. The NOE intensity for R34 is low relative to average backbone values in ancCC-, ancR1- and ancR7-ΔR, suggesting that the guanidinium moiety of R34 displays faster motion than the proteins’ overall tumbling (**Fig. 6b**). The decrease in the relative NOE intensity of R34 between ancCC-ΔR and ancR1-ΔR is consistent with an increase in R34 side chain mobility in ancR1-ΔR. This increased flexibility may be attributable to increased available space in the active site due to the L108V substitution (**Fig. 5c**). The relative NOE intensity increases again in ancR7-ΔR, suggesting that R34 becomes less flexible in this advanced evolutionary intermediate. Foremost, a large change in chemical shift was observed, indicating a change in R34’s average chemical environment (**Fig. 6c**). The modulation in mobility and average location of R34 was recapitulated in MD simulations that included ancR3 (**Fig. 6d**). A significant narrowing in the ensemble of accessible arginine rotamers was observed along the evolutionary trajectory, with ancR1’s ensemble being most dispersed (**Fig. 6d**). The increased mobility of R34 in ancR1 compared to both the starting point, ancCC, and the advanced intermediates, ancR3 and ancR7, was also reflected in the root-mean square fluctuation (RMSF) of its side chain (**Supplementary Fig. 14**). We conclude that the emergence of CHI catalysis occurred via reshaping of the active-site cavity to allow the substrate to bind in a productive mode, thereby also repositioning R34 and enabling it to exert its catalytic effect^20^.

**Figure 6.**
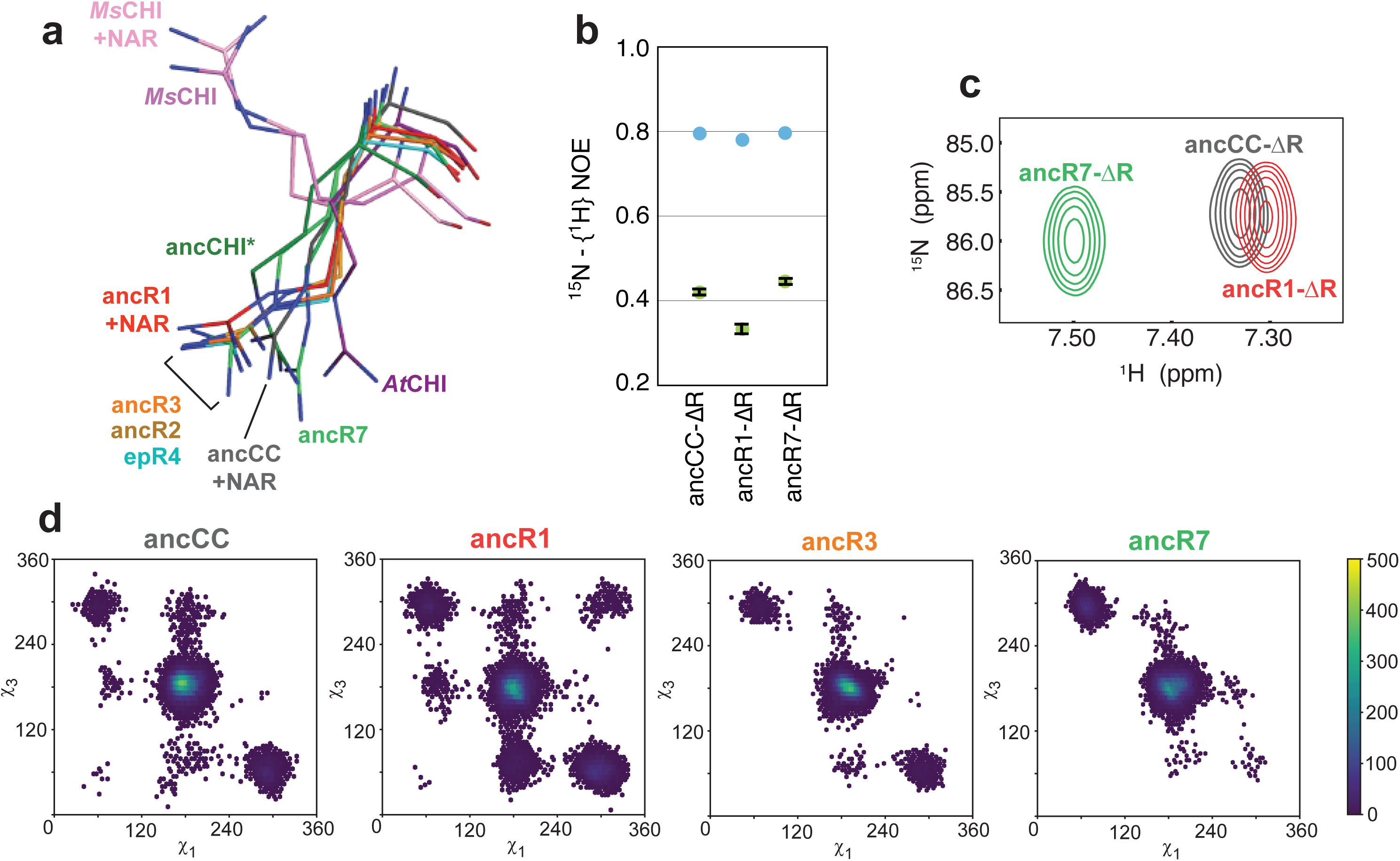
The conformational ensemble of the catalytic Arginine changes over the evolution. (**a**) The structural alignment of different crystal structures reveals that the conformation of R34 varies widely. The presence of (*S*)-naringenin (NAR) does not significantly influence this conformation. The structure of *At*CHI (PDB: 4doi) was reported in [7], the structure of *Ms*CHI with (1eyq) and without (1eyp) NAR was reported in [19]. (**b**) NMR steady-state heteronuclear NOE values (15N-{1H} NOE) for the catalytic arginine, R34. Guanidine 15Nε −1Hε bond vectors (green), and average values for backbone amide 15N-1H bond vectors are shown for ancCC-, ancR1-, and ancR7-ΔR (blue, using resonance peaks downfield of 8.5 ppm). Low 15N-{1H} NOE values indicate mobility on the pico-nanosecond timescale relative to overall molecular tumbling. These experiments reveal that the mobility of the arginine side chain first increases, then decreases at key points in the evolutionary trajectory but remains mobile on the pico-nanosecond timescale for the most active variant, ancR7-ΔR. Average values of triplicate experiments are shown and error bars are standard deviations. (**c**) The distinct HSQC signals of the arginine side chain are indicative of a change in its chemical environment over the evolution. (**d**) Correlated distribution of the χ1 and χ3 dihedral angle values sampled by the Arg34 side chain in ancCC, ancR1, ancR3, and ancR7. Two-dimensional histograms were calculated from the last 50 ns of the simulation of each variant, repeated in five independent replicas, i.e. 250 ns of sampling in each case (or four replicas/200 ns in the case of ancR3), and the angle values are reported in degrees.

## Discussion

Our results indicate that CHI evolved from a catalytically inactive ancestor thus showing that emergence of catalysis and stereospecificity in non-catalytic scaffolds is a feasible evolutionary scenario. That said, a bacterial CHI^22^ has evolved independently in an enzymatic protein fold, exemplifying that plant CHI is an exception rather than the rule. The actual, historical ancestor may have exhibited some isomerase activity, but this activity is unlikely to have exceeded a low, promiscuous level. The three founder mutations that initiate CHI activity were individually inferred in ancCC with tangible probabilities (p(I36)=0.11, p(L99)=0.14, p(L108)=0.04). However, the likelihood that all three were present, giving rise to a relatively active enzyme (**Fig. 3A**; specific activity ∼6×10^3^ µmol/min/mg), is negligible (p<0.001).

That the inferred ancestor is devoid of CHI activity is also interesting because other inferred enzyme ancestors were found to be multi-functional, typically harboring varying levels of the activities found in the extant protein families that diverged from them^23^. Here, we have obtained an ancestor that is not multi-functional, at least with respect to the isomerase activity. However, what the function of this ancestor was remains unknown, foremost because the function of contemporary CHILs is also unknown. Presumably, the function of ancCC or extant CHILs relates to binding of phenylpropanoids as suggested by binding of (*S*)-naringenin to ancCC in a mode that differs from CHI’s.

Unraveling the ancestral function is a challenge also because at the time when ancCC and the early CHI precursors emerged, flavonoid biosynthesis must have been at its infancy^24^. This ancestral context is obviously lost in modern plants. Nonetheless, in the context of an ancient, rudimentary metabolism, emergence from a non-enzymatic protein, and via intermediates that are relatively inefficient enzymes as shown here, seems plausible. Given the high spontaneous rate of chalcone cyclization (half-life for isomerization ∼3.6 h), a weak catalyst or even a binding protein that directs enantioselectivity could prove beneficial to the fitness of the organismal host^25,26^.

CHI’s evolution seems unique not only in having its origins in an inactive ancestor, but also in following a smooth, largely additive trajectory. In fact, we surmise that these two features, and also the relatively facile spontaneous rate of chalcone cyclization, are related; namely, it is the absence of strong epistasis and the pre-existing uncatalyzed cyclization rate that made ancCC a feasible evolutionary starting point despite its lack of initial activity. Multiple parallel trajectories to CHI activity exist – four different founder mutations were identified and various additive combinations thereof (**Fig. 3** and **4**). This trend is in contrast to previous studies supporting the prevalence of epistasis in protein evolution, including sign epistasis, and especially between founder mutations^14,16,27–33^. Beyond enzymatic function, the mutations that initiated CHI’s function affected stability – the apparent melting temperatures of the evolutionary intermediates were reduced by up to 13 °C compared to ancCC (**Supplementary Table 3**). Given the high stability of ancCC^34^, this destabilization had no phenotypic effects, but the actual, historical intermediates may have been affected. Nevertheless, in terms of function, the mutations along CHI’s emergence show uncharacteristically additive effects. This may relate to the preexistence of three key catalytic residues; however, key catalytic residues also preexisted in enzyme trajectories that showed strong epistasis^14,16,35,36^. Low energetic and steric demands of the reaction may be another factor, but these are also not unique to this trajectory^14,16^. A more likely explanation in our view is that the combined likelihood of emergence from a completely inactive starting point and via a rugged, epistatic trajectory is exceedingly low. In other words, for an enzymatically inactive protein to be a viable starting point, founder mutations must arise with relatively high likelihood.

Finally, that CHI’s ancestor is inactive despite all key catalytic residues being in place, analogous to the current limitations of *de novo* enzyme design: That catalytic residues can be placed within a suitable cavity, yet without taking into account dynamics, resulting in geometries that only roughly match catalytically productive conformations, yields poor enzymes at best which generally need to be improved by directed evolution^37–39^. The presence of catalytic residues is necessary, but not sufficient. However, once an initial activity level arise, mutations in other residues, often distal from the active site, enable large improvements in catalytic efficiency^40,41^. As seen in the evolution of CHI, subtle changes in substrate positioning and in the position and flexibility of the catalytic residues relative to the bound substrate can trigger activity in an inactive pocket.

## Acknowledgments

We thank Kesava-Phaneendra Cherukuri for help with the synthesis of chalconaringenin, Brendan Duggan and Xuemei Huang for assistance with NMR, Gordon Louie for assistance with protein x-ray data collection and processing, and George Cortina for help with analyzing the simulations. This work was funded by the Israel Science Foundation Grant 980/14 and the Sasson & Marjorie Peress Philanthropic Fund (D.S.T.); the United States National Science Foundation grant EEC-0813570 (J.P.N.); the Knut and Alice Wallenberg Foundation, Wenner-Gren Foundations and the European Research Council (S.C.L.K.). Computer time was provided by the Swedish National Infrastructure for Computing. J.P.N. is the Arthur and Julie Woodrow Chair and a Howard Hughes Medical Institute investigator. D.S.T. is the Nella and Leon Benoziyo Professor of Biochemistry.

## Author Contributions

M.K. performed ancestral inference with assistance from A.R. M.K. performed directed evolution. M.K., M.D., and J.R.B. performed mutagenesis, protein expression, stable isotope labeling, and biochemical characterization of the proteins. J.R.B., M.K., D.S.T. and J.P.N. performed and analyzed protein x-ray crystallography and NMR. A.P. and S.C.L.K. performed and analyzed MD simulations with assistance from F.S.M. D.S.T. and J.P.N. planned and directed the project, and, together with M.K., J.R.B., A.P., and S.C.L.K., designed the experiments. M.K., J.R.B., J.P.N. and D.S.T. wrote and edited the manuscript.

## Online Methods

### Ancestral reconstruction

To generate a phylogenetic tree, we mined the 1KP transcriptome database^42^ for CHI and CHIL sequences. We also included sequences of several biochemically characterized CHI proteins as well as sequences from ncbi protein BLAST. We covered all major plant lineages (liverworts, mosses, lycophytes, ferns, gymnosperms, and angiosperms) except for hornworts where no CHI or CHIL sequences could be identified among the three transcriptomes (and zero genomes) available at the time. While mosses contain CHILs, no CHI sequences were found in this lineage. Due to the large number of available transcriptomes, redundant sequences (≥ 70%) were removed. However, because the early taxa (liverworts, mosses) were underrepresented in the databases, we included several such sequences despite higher identity. Overall, 88 CHI and 43 CHIL sequences were included (**Supplementary Table 1**). Six previously reported FAPb sequences that are the most closely related to CHI and CHIL^7^ were included as outgroup. A structure-based alignment (**Supplementary Fig. 1**, **Supplementary Data Set 1**) was created with Expresso^43^ using the crystal structures of *At*CHI (PDB ID 4DOI, TAIR gene accession no. At3g55120), CHIL (4DOK, At5g05270), and FAPb (4DOL, At1g53520). Poorly aligned regions (loops, N- and C-termini) were trimmed. Positions that could not be trimmed unambiguously were initially kept. A phylogenetic tree was generated with MrBayes^44^ and showed a consensus from a 1 million generation run (**Fig. 1b**; **Supplementary Fig. 2**). The tree was largely consistent with the tree of life. Note that moss CHILs take up the outgroup position within the CHIL clade. However, the relationship between basal land plants is still under debate^45^. AncCHI, ancCHIL, and ancCC were predicted using FastML and the JTT substitution matrix^46^. Alternative substitution models were tested and gave nearly identical results. By default, we chose the amino acids with the highest posterior probabilities to derive the most probable ancestral sequences. Remaining gaps and ambiguously aligned positions were handled manually as described in detail in **Supplementary Fig. 3** and **Supplementary Table 2**. The N- and C-terminal amino acids were predicted with large ambiguity and hence, identical adaptor sequences derived from *At*CHI were added to each ancestor (**Supplementary Fig. 1**, **Supplementary Table 2**). Ultimately, we obtained three 220-amino acid proteins, with pairwise sequence identities of 89% for ancCC/ancCHIL, 70% for ancCC/ancCHI, and 60% for ancCHI/ancCHIL.

### Protein expression and purification

Synthetic genes for the three ancestors were ordered from Hylabs. All variants were cloned into pHis8 (a modified pET28 vector containing an N-terminal His8-tag) using Nco I and Hind III. Plasmids were transformed into *E. coli* BL21 (DE3). Cells were grown at 37 °C in LB containing 50 µg/mL Kanamycin to an OD OD_600nm_ of 0.6-1. Overnight expression was induced by addition of 1M IPTG and the temperature reduced to 18 °C. Cells were harvested by centrifugation, resuspended and lysed by sonication in Column Washing buffer (50 mM Tris-HCl, pH 8.0, 500 mM NaCl, 20 mM imidazole, 10% glycerol) supplemented with 1% Tween 20, 10 mM β-mercaptoethanol, 100 µg/mL Lysozyme, benzonase (Novagen) at a dilution of 10^4^-10^5^, and protease inhibitor cocktail for purification of His-tagged proteins (Sigma). Cell debris was removed by centrifugation and the clarified lysate passed through a 45 µM filter prior to purification with Nickel beads (Adar Biotech). The Elution buffer differed from the Column Washing buffer in that it contained 250 mM imidazole. Proteins were dialyzed overnight against Activity buffer (50 mM HEPES pH 7.5, 150 mM NaCl) and concentrated over spin-column concentrators if necessary. Protein concentration was determined using the Pierce BCA Protein Assay Kit (Thermo Fisher Scientific).

### Synthesis of chalconaringenin

Chalconaringenin was synthesized as previously described^47^ with minor modifications. Naringenin (2 g, Sigma-Aldrich) was treated with 5% NaOH in methanol (50 mL) under reflux for 2 hours. Subsequently, 200 mL of 1 M HCl were added to the mixture and the aqueous layer was extracted three times with ethyl acetate. The combined organic layers were washed with brine and water, dried over Na_2_SO_4_, and evaporated. The crude product was further purified by silica gel column chromatography using a hexane:ethylacetate gradient (100% hexane, then 7:1, 3:1, 2:1). ^1^H NMR (400 MHz, DMSO-*d*_*6*_): δ 12.56 (s, OH-2’/6’), 10.46 (s, OH-4’), 10.12 (s, OH-4), 7.95 (d, *J*=15.57 Hz, H-β), 7.64 (d, *J*=15.57 Hz, H-α), 7.51 (d, *J*=8.32 Hz, H-2/6), 6.81 (d, *J*=8.32 Hz, H- 3/5), 5.83(s, H-3’/5’); ^13^C NMR (100 MHz, DMSO-*d*_*6*_): δ 191.88, 164.86, 164.54, 159.96, 142.49, 130.50, 126.26, 123.96, 116.14, 104.39, 95.03.

### Enzyme assays

Naringenin formation was followed by the decrease in absorbance at 390 nm (chalconaringenin is yellow; naringenin is colorless), typically in a Synergy HT spectrophotometer (Bio-TEK). To determine the activity in crude cell lysates, cells were grown at a 500 µL culture volume in 96-well plates and lysed as described below under *Screening*. Lysates were diluted (from no dilution up to a 10,000-fold dilution for extant CHIs) to determine initial rates of chalconaringenin isomerization (in the range of µM/min). One volume of 200 µM chalconaringenin in 50 mM HEPES, pH 7.5 and 5% ethanol was added to one volume of diluted cell lysate. Initial rates were normalized to cell density (OD_600nm_), corrected for the dilution factor, and averaged. For the data shown in **Fig.s 2–4** and **Tables S4** and **9**, each variant was grown in at least two wells and the average determined. The whole experiment (cell growth, protein expression, lysis and enzymatic assay) was repeated three times and the combined average determined. For directed evolution, each well contained a different variant (except for the respective control, *i.e.* the best variant from the previous round, which was grown in at least triplicate on each plate). Improved variants were regrown and assayed in triplicate.

Specific activity was determined with purified proteins (µmol product generated per min per mg protein) at a substrate concentration of 100 µM. For determination of kinetic parameters, assays were performed at 12 different substrate concentrations (4.5 – 180 µM) in 50 mM HEPES pH 7.5 with 5% ethanol as co-solvent. The spontaneous, background rates in buffer were subtracted, and the net initial rates were directly fit to the Michaelis-Menten model using Kaleidagraph. In cases where no significant rate saturation was observed, *k*_*cat*_/*K*_*M*_ was extracted by a linear fit. The rate constant of the spontaneous reaction was calculated from a linear fit of initial rates in buffer (*k*_*unca*t_ = 1.05 × 10^-3^s^-1^, **Supplementary Fig. 5**). The spontaneous isomerisation rate of chalconaringenin is relatively high. Hence, for the early evolutionary intermediates, assays were performed at enzyme concentration that exceeded substrate concentration ([E]_0_ > [S]_0_). These could reveal k_cat_ values that are lower than k_uncat_. For those proteins reported as inactive (*e.g.* ancCC) no activity above the background rate was detected at up to 50 µM protein.

### Determination of enantioselectivity

Proteins were incubated with 60 µM chalconaringenin in 50 mM HEPES pH 7.5 for 30 minutes at room temperature. For low-activity mutants, up to 1 mM of protein (>10-fold excess over substrate) was used. Naringenin was extracted with 0.5 volumes of ethyl acetate. The organic phase was concentrated in a SpeedVac, centrifuged to remove residual precipitated protein, and the sample analyzed on a chiral HPLC column (Lux 5u Cellulose-4, 150 x 4.6 mm, Phenomenex) using a linear gradient (88% hexane/12% ethanol, 0.1% TFA) and a flow rate of 1.5 ml/min. Chalconaringenin (retention time 5.6 min), R-naringenin (8.1 min) and S-naringenin (6.7 min) were detected by absorbance at 310 nm, where substrate and products absorb to a similar extent. The traces of (*R*)-naringenin detected in case of the low-activity variants are likely due to the spontaneous background reaction that generates racemic product.

### Thermostability assays

Solutions of 10 µM protein were heated with 10x SYPRO Orange (Sigma) in 50 mM HEPES pH 7.5 to a final temperature of 25 up to 95 °C, at a ramp of 0.9 °C/min, in a Viia 7 Real-Time PCR system (Applied Biosystems). The increase in fluorescence that likely results from protein unfolding was monitored (excitation 488 nm, emission 500-750 nm). The apparent melting temperature (T_m_) is the midpoint of the resulting temperature-fluorescence transition curve and was determined from the minimum value of the first derivative. Measurements were performed in triplicate and average T_m_ values determined. The stability of selected variants was also measured by following changes in tryptophan fluorescence upon heat-induced unfolding in a Prometheus NT.48 NanoDSF instrument (NanoTemper Technologies). Solutions of 10 µM protein were heated in 50 mM HEPES pH 7.5 from 20 to 95 °C at a ramp of 2 °C/min, the observed fluorescence ratios at (excitation 280 nm; emission 330nm/350nm) were plotted against temperature, and T_m_ values determined as the minimum of the first derivative.

### Cloning and library construction

#### Phylogenetic libraries

To reduce library size, we sought to remove neutral mutations. To this end, an evolutionary conservation analysis was performed with ConSurf^48^ (**Supplementary Fig. 6a**). Considering also the location of each substitution in the protein structure (using the crystal structure of *Medicago sativa* CHI with bound product, PDB ID 1eyq), we identified and reverted 29 non-conserved and/or peripheral substitutions in ancCHI to ancCC’s amino acids, thus yielding ancCHI* (**Supplementary Fig. 6b**). The catalytic activity of ancCHI* was confirmed as similar to ancCHI (∼2-fold lower *k*_*cat*_/*K*_*M*_, **Supplementary Table 4**). The libraries were created with ISOR (Incorporating Synthetic Oligonucleotides via Gene Reassembly^49^) at an average mutation rate of 2 amino acid exchanges/gene. For the round 1 library, ancCC was shuffled with 39 oligonucleotides, each containing one mutation to ancCHI’s amino acid (**Supplementary Table 5**). In cases where the amino acid exchange could only have occurred by two base substitutions, the oligonucleotide was partially randomized to include the plausible intermediary amino acid(s). Because several of the ancestral exchanges were at nearby positions, the primer set was updated after each round to avoid the reversion of ancCHI mutations that were fixated in the previous rounds (see **Supplementary Table 5** for an example). Overall, seven rounds of library construction and screening were performed. Low-mutation rate libraries were generated in the same manner, but at a mutation rate of 0.2 amino acid exchanges/gene. To ensure that all single mutations were represented with significant probability, ∼930 variants were screened for each of the four low-rate libraries (∼5× oversampling). More details on library generation and screening are given in **Supplementary Table 6**.

#### Random mutagenesis libraries

Random libraries were created by alternating rounds of error-prone PCR and DNA shuffling. Error-prone PCR was performed using the GeneMorph II Random Mutagenesis kit (Agilent) according to the manufacturer’s instructions. The average mutation rate was determined by sequencing a sample of randomly selected variants, and libraries with an average of 1.2 amino acid substitutions/gene (∼2 nucleotide substitutions/gene) were screened. DNA shuffling was performed with StEP (Staggered Extension Process)^50^. More details on library generation and screening are given in **Supplementary Table 6**.

### Site-directed mutagenesis

Mutants were constructed as described in the QuikChange Site-Directed Mutagenesis manual (Agilent).

### Library screening

Libraries were clones into pHis8, transformed into *E. coli* BL21 (DE3) and plated on kanamycin LB-agar plates. Single colonies were picked into deep 96-well plates and grown overnight at 30 °C in 200 µL LB per well supplemented with 50 µg/mL kanamycin. Wells containing fresh LB/Kanamycin (500 µL) were inoculated with 25 µL of this preculture and protein expression was induced as above. Cells pelleted by centrifugation and the supernatant removed. Cell lysis was achieved by freezing for 30 minutes at −80 °C, resuspension in 200 µL of Activity buffer supplemented with 0.1% Triton-X100, 100 µg/mL Lysozyme, and 0.25-2.5 U/mL benzonase (Novagen) and incubation for 30 minutes at room temperature. Cell debris was pelleted by centrifugation and 20-100 µL of clarified lysate (or dilutions thereof, depending on the activity level of the library) were removed to assay for CHI activity. Substrate buffer (Activity buffer supplemented with 1% DMSO and chalconaringenin) was added to a final volume of 200 µL at a final substrate concentration of 50-180 µM, and the initial rates of naringenin formation were determined. Improved variants (compared to the best variant of the previous round) were picked and re-grown in triplicate, initial rates re-measured, and the average values determined. Plasmid DNA was isolated and variants were sequenced (**Supplementary Tables 7, 10** and **11**). The plasmids were retransformed, and the resulting single colonies were used to inoculate new cultures for activity determination and protein purification.

### X-ray crystallography

Proteins purified by Ni-NTA affinity chromatography were prepared for crystallization by elution from a Superdex 200 size exclusion column in buffer containing 25 mM Tris (pH 8.0), 200 mM NaCl and 2 mM DTT. Proteins were crystallized by hanging drop vapor diffusion at 4 °C. Protein crystals of ancCC and ancR1 grew overnight at a concentration of 150 mg/ml in 42% MPD and 50 mM acetic acid (pH 4.5). Crystals were then transferred to a condition with 65% MPD, 50 mM acetic acid (pH 4.5), 20 mM naringenin and soaked for three days. Protein crystals of ancR2 were grown overnight at a concentration of 100 mgs/ml in 25% PEG 4K, 2.5 M ammonium formate and 100 mM succinic acid (pH 5.5). Protein crystals of ancR5 were grown over 2 weeks at a concentration of 140mgs/ml in 3.5mM ammonium sulfate and 100mM TAPS (pH 8.5). Protein crystals of ancR7 were grew over one month at a concentration of 50 mgs/ml in 500mM NaCl, 100mM NaCitrate tribasic dihydrate (pH 5.6), and 2% ethylene imine polymer. Protein crystals of ancR3 formed overnight at a concentration of 200 mg/ml in 25% PEG 4K, 2.5 M ammonium formate and 100 mM succinic acid, pH 5.5. Protein crystals of ancCHI* grew over one week at a concentration of 100mg/ml in 4.5 M NaCl and acetic acid, pH 4.5. Protein crystals of epR4 grew in two days at a concentration of 120 mg/ml in 29% PEG 4K, 2.5 M ammonium formate and 50 mM succinic acid, pH 5.5. Protein crystals were frozen in liquid nitrogen and data were collected at 100 K at beamlines 8.2.1 and 8.2.2 at the Advanced Light Source, Lawrence Berkeley National Laboratory. Data were integrated with iMOSFLM^51^ and scaled with SCALA^52^. *Arabidopsis thaliana* CHI (*At*CHI, PDB: 4DOI) was used as a search model for molecular replacement^7^, and phases were solved with PHASER^53^. Models were modified in COOT^54^ and refined with PHENIX^55^. Statistics for data collection and refinement are in **Supplementary Table 14**. Coordinates and structure factors have been deposited in the Protein Data Bank under accession codes 5WKR (ancCC), 5WKS (ancR1), 5WL3 (ancR2), 5WL4 (ancR3), 5WL5 (ancR5), 5WL6 (ancR7), 5WL7 (ancCHI*), and 5WL8 (epR4).

### NMR

Constructs of ancCC, ancR1 and ancR7 were generated with the substitutions: R115K, R117K, and R136K for the purpose of unambiguously resolving the ^1^H^ε^ –^15^N^ε^ arginine signal of R34. The specific activity of ancR1-ΔR was essentially the same compared to ancR1 (1.7-fold reduction). While the specific activity of ancR7-ΔR was 15-fold lower than that of its parent ancR7, it is still a highly active enzyme (**Supplementary Table 9**). AncCC-ΔR, like ancCC, shows no activity, but is well-folded as shown by the NMR data and therefore likely functional. Proteins were expressed using ^15^N isotopically-labeled ammonium sulfate in minimal media, and purified by affinity chromatography followed by removal of the His8-tag and finally ion exchange chromatography. The protein samples were then prepared at a final concentration of 1mM in a buffer composed of 50 mM NaPO_4_, 5 mM dithiothreitol, and 10% D_2_O (pH 6.1). ^1^H–^15^N HSQC spectra were measured at 298 K on an 800 MHz Bruker AVANCE NEO solution state NMR spectrometer, equipped with a 5 mm cryogenically cooled TCI probe. ^15^N–{^1^H}NOEs were measured with presaturated and unsaturated experiments and recorded interleaved with a relaxation delay of 5.5 seconds. ^15^N– {^1^H}NOE values were calculated from the ratio of peak intensities from the presaturated to the unsaturated experiments. Experiments were performed in triplicate to estimate error based on standard deviation between the calculated ^15^N– {^1^H}NOE value for each separate run. Unassigned amide backbone peaks downfield of 8.5 ppm represent structured residues in the protein and were selected to calculate the average heteronuclear NOE of backbone values for comparison with the R34 sidechain. ^15^N longitudinal (T1) relaxation experiments were performed at 800 MHz with 2 duplicate points for error estimation. Experiments were collected with T1 delays of 30, 300, 600(x2), 900, 1200(x2), 1800, 2400, and 3600ms. Peak intensities were measured in Sparky^56^ and fitting of the exponential decay curves was performed using the RelaxGUI^57^. Calculated T1 values for the R34 ^1^H^ε^ –^15^N^ε^ resonance are 1.137 ± 0.029 sec, 1.154 ± 0.029 sec, and 0.9691 ± 0.035 sec for ancCC(3K), ancR1(3K) and ancR7(3K), respectively. Data were processed using NMRpipe^58^, Sparky^56^ and Relax^57^.

### MD Simulations

Simulations of both the substrate-free enzymes, as well as substrate-bound complexes were performed on ancCC, ancR1, ancR3, ancR7, and additionally on ancCHI and *At*CHI for the enzyme-substrate complexes, starting from the coordinates provided in the respective PDB IDs listed under “X-ray crystallography” and 4DOI for *At*CHI^7^. Only Chain A was retained for the simulations, and all heteroatoms and water molecules further than 4 Å away from the protein were removed. Missing N- and C-terminal residues were modeled with Modeller v. 9.18^59^, using the standard presets, and selecting coordinates based on the lowest z-score. The ionization states of all titratable residues, and the protonation states of histidine side chains, was determined by empirical screening with PROPKA^60^ and histidine protonation patterns were assigned based on visual examination of the local environment of each side chain. The substrate was placed manually in the active site by analogy with *Ms*CHI (PDB ID 1EYQ^19^) for the productive (CHI type II like) binding mode, and the ancR1 structure for the non-productive binding mode, respectively. The only exception to this was ancCC, where a structure with the product in the non-productive binding mode was available to use as a template. In addition to the simulations of the substrate-bound complexes, simulations of ancCC and ancR1 variants in complex with the product were performed. These simulation were based on the crystallographic coordinates of the two systems with the product bound (PDB IDs 5WKR and 5WKS), and were performed in order to validate that the simulation protocol can reproduce the stability of the product complex found in the non-productive binding mode in those crystal structures. The model systems for these simulations were prepared using the same procedure as described above for the simulations of substrate-free enzymes and substrate-bound complexes.

All simulations were performed using the AMBER16 simulation package^61^ using the AMBER ff14SB force field^62^ and the General Amber Force Field (GAFF)^63^ for the protein and substrate/product, respectively. Partial charges for chalconaringenin were calculated at the HF/6-31G* level of theory, using the Gaussian09 simulation package^64^, and fitted using the standard restrained electrostatic potential (RESP) procedure^65^ (**Supplementary Table 16**). All starting structures for the simulations were prepared using the LEaP module of AMBER. The structures were solvated in a truncated octahedron of TIP3P^66^ water molecules, extending to at least 10Å from the protein atoms. Na^+^ and Cl^-^ ions were added to obtain a charge neutralized system with an ion concentration of 0.15 M.

All systems were initially minimized (1000 steps steepest descent followed by conjugate gradient minimization) with a 25 kcal mol^-1^ Å^-2^ harmonic positional restraint applied to all solute atoms, followed by two more minimization steps with the restraint dropped to 5 kcal mol^-1^ Å^-2^. The minimized systems were then gradually heated from 100 to 300K over 40 ps simulation time at constant volume (NVT ensemble), with the temperature equilibrated for a further 60 ps. The density of the system was then equilibrated by performing 300 ps of constant pressure (NPT) simulations with isotropic position scaling, using the Berendsen barostat^67^ for pressure control. The 5 kcal mol^-1^ Å^-2^ positional restraint was retained for both heating and density equilibration, and this restraint was gradually released over 350 ps of NVT simulations, with the restraint completely removed for a final 50 ps of equilibration time. An additional 50 kcal mol^-1^ rad^2^ harmonic restraint was applied to the C1-C2-C3-C4 torsion angle of the substrate (see **Supplementary Table 16** for atom numbering) to keep this torsion angle within the range of −20° to 90°; this restraint was also removed in the last 50 ps of the initial equilibration. Finally, the initially equilibrated systems were subjected to 100 ns of fully unrestrained production simulations in an NVT ensemble. This protocol was repeated five times for each system to generate five independent trajectories, resulting in 500 ns of simulation time per system, the RMSD convergence of which is shown in **Supplementary Fig. 15**. A 2 fs time step was used throughout, and all bonds involving hydrogen atoms were constrained using the SHAKE algorithm^68^. The temperature was controlled using Langevin dynamics^69^ with a 1 ps^-1^ collision frequency, and long-range electrostatic interactions were calculated using the Particle Mesh Ewald (PME) method^70^. An 8Å cutoff was used to describe non-bonded interactions.

Finally, all simulations were analyzed using CPPTRAJ from the AmberTools package^61^, Visual Molecular Dynamic package^71^ and MDTraj 1.8.0 library^72^, and the structure graphics were prepared using UCSF Chimera package^73^.

## References

1. Adrain, C. & Freeman, M., New lives for old: evolution of pseudoenzyme function illustrated by iRhoms. Nat Rev Mol Cell Biol 13, 489–498 (2012).

2. Ortmayer, M. et al., An oxidative N-demethylase reveals PAS transition from 676 ubiquitous sensor to enzyme. Nature 539, 593–597 (2016).

3. Taga, M.E. et al., BluB cannibalizes flavin to form the lower ligand of vitamin B12. Nature 446, 449–453 (2007).

4. Tam, R. & Saier, M.H., Jr., A bacterial periplasmic receptor homologue with catalytic activity: cyclohexadienyl dehydratase of Pseudomonas aeruginosa is homologous to receptors specific for polar amino acids. Res Microbiol 144, 165–169 (1993).

5. Yuhara, K. et al., Enzymatic characterization and gene identification of aconitate isomerase, an enzyme involved in assimilation of trans-aconitic acid, from Pseudomonas sp. WU-0701. FEBS J 282, 4257–4267 (2015).

6. Koes, R.E., Quattrocchio, F., & Mol, J.N.M., The Flavonoid Biosynthetic Pathway in Plants: Function and Evolution. BioEssays 16, 123–132 (1994).

7. Ngaki, M.N. et al., Evolution of the chalcone-isomerase fold from fatty-acid binding to stereospecific catalysis. Nature 485, 530–533 (2012).

8. Morita, Y. et al., A chalcone isomerase-like protein enhances flavonoid production and flower pigmentation. Plant J 78, 294–304 (2014).

9. Jiang, W. et al., Role of a chalcone isomerase-like protein in flavonoid biosynthesis in Arabidopsis thaliana. J Exp Bot 66, 7165–7179 (2015).

10. Jez, J.M., Bowman, M.E., & Noel, J.P., Role of hydrogen bonds in the reaction mechanism of chalcone isomerase. Biochemistry 41, 5168–5176 (2002).

11. Bar-Rogovsky, H. et al., Assessing the prediction fidelity of ancestral reconstruction by a library approach. Protein Eng Des Sel 28, 507–518 (2015).

12. Eick, G.N. et al., Robustness of Reconstructed Ancestral Protein Functions to Statistical Uncertainty. Mol Biol Evol 34, 247–261 (2017).

13. Randall, R.N. et al., An experimental phylogeny to benchmark ancestral sequence reconstruction. Nat Commun 7, 12847 (2016).

14. Tokuriki, N. et al., Diminishing returns and tradeoffs constrain the laboratory optimization of an enzyme. Nat Commun 3, 1257 (2012).

15. Tokuriki, N. et al., The stability effects of protein mutations appear to be universally distributed. J Mol Biol 369, 1318–1332 (2007).

16. Salverda, M.L. et al., Initial mutations direct alternative pathways of protein evolution. PLoS Genet 7, e1001321 (2011).

17. Kaltenbach, M. et al., Reverse evolution leads to genotypic incompatibility despite functional and active site convergence. Elife 4 (2015).

18. Dickinson, B.C. et al., Experimental interrogation of the path dependence and stochasticity of protein evolution using phage-assisted continuous evolution. Proc Natl Acad Sci U S A 110, 9007–9012 (2013).

19. Jez, J.M., Bowman, M.E., Dixon, R.A., & Noel, J.P., Structure and mechanism of the evolutionarily unique plant enzyme chalcone isomerase. Nat Struct Biol 7, 786–715 (2000).

20. Burke, J.R. et al., Origin of Enantioselective Catalysis by a Guanidine Moiety in Chalcone Isomerases. submitted manuscript (2017).

21. Farrow, N.A. et al., Backbone dynamics of a free and phosphopeptide-719! complexed Src homology 2 domain studied by 15N NMR relaxation. Biochemistry 33, 5984–6003 (1994).

22. Thomsen, M. et al., Structure and catalytic mechanism of the evolutionarily unique bacterial chalcone isomerase. Acta Crystallogr D Biol Crystallogr 71, 907–917 (2015).

23. Gumulya, Y. & Gillam, E.M., Exploring the past and the future of protein evolution with ancestral sequence reconstruction: the 'retro' approach to protein engineering. Biochem J 474, 1–19 (2017).

24. Weng, J.K. & Chapple, C., The origin and evolution of lignin biosynthesis. New Phytol 187, 273–285 (2010).

25. Bar-Even, A. & Salah Tawfik, D., Engineering specialized metabolic pathways-is there a room for enzyme improvements? Curr Opin Biotechnol 24, 310–319 (2013).

26. Keller, M.A., Piedrafita, G., & Ralser, M., The widespread role of non-732! enzymatic reactions in cellular metabolism. Curr Opin Biotechnol 34, 153–161 (2015).

27. Breen, M.S. et al., Epistasis as the primary factor in molecular evolution. Nature 490, 535–538 (2012).

28. de Visser, J.A., Cooper, T.F., & Elena, S.F., The causes of epistasis. Proc Biol Sci 278, 3617–3624 (2011).

29. Harms, M.J. & Thornton, J.W., Evolutionary biochemistry: revealing the historical and physical causes of protein properties. Nat Rev Genet 14, 559–571 (2013).

30. Kaltenbach, M. & Tokuriki, N., Dynamics and constraints of enzyme evolution. J Exp Zool B Mol Dev Evol 322, 468–487 (2014).

31. McCandlish, D.M. et al., The role of epistasis in protein evolution. Nature 497, E1–2; discussion E2-3 (2013).

32. Poelwijk, F.J., Kiviet, D.J., Weinreich, D.M., & Tans, S.J., Empirical fitness landscapes reveal accessible evolutionary paths. Nature 445, 383–386 (2007).

33. Whitlock, M.C., Phillips, P.C., Moore, F.B.-G., & Tonsor, S.J., Multiple Fitness Peaks and Epistasis. Annu. Rev. Ecol. Syst. 26, 601–629 (1995).

34. Trudeau, D.L., Kaltenbach, M., & Tawfik, D.S., On the Potential Origins of the High Stability of Reconstructed Ancestral Proteins. Mol Biol Evol (2016).

35. Noor, S. et al., Intramolecular epistasis and the evolution of a new enzymatic function. PLoS One 7, e39822 (2012).

36. Lozovsky, E.R. et al., Stepwise acquisition of pyrimethamine resistance in the malaria parasite. Proc Natl Acad Sci U S A 106, 12025–12030 (2009).

37. Kiss, G. et al., Computational enzyme design. Angew Chem Int Ed Engl 52, 5700–5725 (2013).

38. Kries, H., Blomberg, R., & Hilvert, D., De novo enzymes by computational design. Curr Opin Chem Biol 17, 221–228 (2013).

39. Lassila, J.K., Conformational diversity and computational enzyme design. Curr Opin Chem Biol 14, 676–682 (2010).

40. Blomberg, R. et al., Precision is essential for efficient catalysis in an evolved Kemp eliminase. Nature 503, 418–421 (2013).

41. Khersonsky, O. et al., Bridging the gaps in design methodologies by evolutionary optimization of the stability and proficiency of designed Kemp eliminase KE59. Proc Natl Acad Sci U S A 109, 10358–10363 (2012).

42. Matasci, N. et al., Data access for the 1,000 Plants (1KP) project. Gigascience 3, 17 (2014).

43. Armougom, F. et al., Expresso: automatic incorporation of structural information in multiple sequence alignments using 3D-Coffee. Nucleic Acids Res 34, W604–608 (2006).

44. Ronquist, F. & Huelsenbeck, J.P., MrBayes 3: Bayesian phylogenetic inference under mixed models. Bioinformatics 19, 1572–1574 (2003).

45. Cronk, Q.C., Plant evolution and development in a post-genomic context. Nat Rev Genet 2, 607–619 (2001).

46. Ashkenazy, H. et al., FastML: a web server for probabilistic reconstruction of ancestral sequences. Nucleic Acids Res 40, W580–584 (2012).

47. Miranda, C.L. et al., Antioxidant and prooxidant actions of prenylated and nonprenylated chalcones and flavanones in vitro. J Agric Food Chem 48, 3876–3884 (2000).

48. Ashkenazy, H. et al., ConSurf 2010: calculating evolutionary conservation in sequence and structure of proteins and nucleic acids. Nucleic Acids Res 38, W529–533 (2010).

49. Herman, A. & Tawfik, D.S., Incorporating Synthetic Oligonucleotides via Gene Reassembly (ISOR): a versatile tool for generating targeted libraries. Protein Eng Des Sel 20, 219–226 (2007).

50. Zhao, H. et al., Molecular evolution by staggered extension process (StEP) in vitro recombination. Nat Biotechnol 16, 258–261 (1998).

51. Battye, T.G. et al., iMOSFLM: a new graphical interface for diffraction-image processing with MOSFLM. Acta Crystallogr D Biol Crystallogr 67, 271–281 (2011). Crystallogr 62, 72–82 (2006).

52. Evans, P., Scaling and assessment of data quality. Acta Crystallogr D Biol Crystallogr 62, 72–82 (2006).

53. McCoy, A.J. et al., Phaser crystallographic software. J Appl Crystallogr 40, 658–791 (2007).

54. Emsley, P., Lohkamp, B., Scott, W.G., & Cowtan, K., Features and development of Coot. Acta Crystallogr D Biol Crystallogr 66, 486–501 (2010).

55. Adams, P.D. et al., PHENIX: a comprehensive Python-based system for macromolecular structure solution. Acta Crystallogr D Biol Crystallogr 66, 213–221 (2010).

56. Goddard, T.D. & Kneller, D.G., SPARKY 3. University of California, San Francisco (2008).

57. Bieri, M., d'Auvergne, E.J., & Gooley, P.R., relaxGUI: a new software for fast and simple NMR relaxation data analysis and calculation of ps-ns and mus motion of proteins. J Biomol NMR 50, 147–155 (2011).

58. Delaglio, F. et al., NMRPipe: a multidimensional spectral processing system based on UNIX pipes. J Biomol NMR 6, 277–293 (1995).

59. Fiser, A. & Sali, A., Modeller: generation and refinement of homology-based protein structure models. Methods Enzymol 374, 461–491 (2003).

60. Sondergaard, C.R., Olsson, M.H., Rostkowski, M., & Jensen, J.H., Improved Treatment of Ligands and Coupling Effects in Empirical Calculation and Rationalization of pKa Values. J Chem Theory Comput 7, 2284–2295 (2011).

61. Maier, J.A. et al., ff14SB: Improving the Accuracy of Protein Side Chain and Backbone Parameters from ff99SB. J Chem Theory Comput 11, 3696–3713 (2015).

62. Case, D.A. et al., AMBER 2016. University of California, San Francisco (2016).

63. Wang, J. et al., Development and testing of a general amber force field. J Comput Chem 25, 1157–1174 (2004).

64. M. J. Frisch, G.W.T., H. B. Schlegel, G. E. Scuseria, M. A. Robb, J. R. Cheeseman, G. Scalmani, V. Barone, G. A. Petersson, H. Nakatsuji, X. Li, M. Caricato, A. Marenich, J. Bloino, B. G. Janesko, R. Gomperts, B. Mennucci, H. P. Hratchian, J. V. Ortiz, A. F. Izmaylov, J. L. Sonnenberg, D. Williams-Young, F. Ding, F. Lipparini, F. Egidi, J. Goings, B. Peng, A. Petrone, T. Henderson, D. Ranasinghe, V. G. Zakrzewski, J. Gao, N. Rega, G. Zheng, W. Liang, M. Hada, M. Ehara, K. Toyota, R. Fukuda, J. Hasegawa, M. Ishida, T. Nakajima, Y. Honda, O. Kitao, H. Nakai, T. Vreven, K. Throssell, J. A. Montgomery, Jr., J. E. Peralta, F. Ogliaro, M. Bearpark, J. J. Heyd, E. Brothers, K. N. Kudin, V. N. Staroverov, T. Keith, R. Kobayashi, J. Normand, K. Raghavachari, A. Rendell, J. C. Burant, S. S. Iyengar, J. Tomasi, M. Cossi, J. M. Millam, M. Klene, C. Adamo, R. Cammi, J. W. Ochterski, R. L. Martin, K. Morokuma, O. Farkas, J. B. Foresman, D. J. Fox, Gaussian 09, Revision D.01. Gaussian, Inc., Wallingford CT (2016).

65. Cieplak, P., Cornell, W.D., Bayly, C., & Kollman, P.A., Application of the multimolecule and multiconformational RESP methodology to biopolymers: Charge derivation for DNA, RNA, and proteins. J Comput Chem 16, 1357–1377 (1995).

66. Jorgensen, W.L. et al., Comparison of simple potential functions for simulating liquid water. The Journal of Chemical Physics 79, 926–935 (1983).

67. Berendsen, H.J.C. et al., Molecular dynamics with coupling to an external bath. Journal of Chemical Physics 81, 3684–3690 (1984).

68. Ryckaert, J.-P., Ciccotti, G., & Berendsen, H.J.C., Numerical integration of the Cartesian Equations of Motion of a System with Constraints: Molecular Dynamics of n-Alkanes. Journal of Computational Physics 23, 327–341 (1977).

69. Feller, S.E., Zhang, Y., Pastor, R.W., & Brooks, B.R., Constant pressure molecular dynamics simulation: The Langevin piston method. The Journal of Chemical Physics 103, 4613–4621 (1995).

70. Darden, T., York, D., & Pederson, L., Particle Mesh Ewald: An N•log(N) Method for Ewald Sums in Large Systems. Journal of Chemical Physics 98, 10089–10092 (1993).

71. Humphrey, W., Dalke, A., & Schulten, K., VMD: visual molecular dynamics. J Mol Graph 14, 33–38, 27-38 (1996).

72. McGibbon, R.T. et al., MDTraj: A Modern Open Library for the Analysis of Molecular Dynamics Trajectories. Biophys J 109, 1528–1532 (2015).

73. Pettersen, E.F. et al., UCSF Chimera-a visualization system for exploratory research and analysis. J Comput Chem 25, 1605–1612 (2004).

